# Effect of feedback regulation on stem cell fractions in tissues and tumors: understanding chemo-resistance in bladder cancer

**DOI:** 10.1101/818005

**Authors:** Lora D. Weiss, P. van den Driessche, John S. Lowengrub, Dominik Wodarz, Natalia L. Komarova

## Abstract

While resistance mutations are often implicated in the failure of cancer therapy, lack of response also occurs without such mutants. In bladder cancer mouse xenografts, repeated chemotherapy cycles have resulted in cancer stem cell (CSC) enrichment, and consequent loss of therapy response due to the reduced susceptibility of CSCs to drugs. A particular feedback loop present in the xenografts has been shown to promote CSC enrichment in this system. Yet, many other regulatory loops might also be operational and might promote CSC enrichment. Their identification is central to improving therapy response. Here, we perform a comprehensive mathematical analysis to define what types of regulatory feedback loops can and cannot contribute to CSC enrichment, providing guidance to the experimental identification of feedback molecules. We derive a formula that reveals whether or not the cell population experiences CSC enrichment over time, based on the properties of the feedback. We find that negative feedback on the CSC division rate or positive feedback on differentiated cell death rate can lead to CSC enrichment. Further, the feedback mediators that achieve CSC enrichment can be secreted by either CSCs or by more differentiated cells. The extent of enrichment is determined by the CSC death rate, the CSC self-renewal probability, and by feedback strength. Defining these general characteristics of feedback loops can guide the experimental screening for and identification of feedback mediators that can promote CSC enrichment in bladder cancer and potentially other tumors. This can help understand and overcome the phenomenon of CSC-based therapy resistance.

## Introduction

Tumor stem cells are thought to be important for the initiation and maintenance of cancers. In addition, it is becoming clear that tumor stem cells can also contribute to a reduced response to treatments, especially if the fraction of the stem cells in the tumor is high [1]. Cancer stem cells (CSCs) are intrinsically less responsive to drug treatment [2-4] due to protective properties they share with normal stem cells, including higher expression of drug-efflux pumps [1], better DNA-repair capacity [5], and enhanced protection against reactive oxygen species [6]. High CSC fractions can therefore give rise to poor a response to therapy even in the absence of resistance-inducing mutations.

CSC-based resistance falls in the more general category of non-genetic drug resistance. There is strong evidence that tumors contain non-genetic, heritable variation, and that this contributes to clonal evolutionary processes [7]. The importance of the interplay between genetic and non-genetic heterogeneity for clonal evolutionary processes has been demonstrated with mathematical models, especially in the context of therapeutic interventions [8,9].

The importance of stem cell-based resistance *in vivo* has been established in patient-derived bladder cancer mouse xenografts [10]. In clinically realistic chemotherapy regimes, it has been demonstrated that stem cell fractions increased during successive treatment cycles, and that this increase in the stem cell fractions correlated with a reduced response to chemotherapy during the next therapy cycle. Experiments identified a positive feedback loop that operates within the tumor as an important component of stem cell enrichment during the treatment phase [10]. Chemotherapy-induced death of more differentiated cells resulted in a wound healing type response that is mediated by PGE2 and that resulted in the activation and proliferation of CSCs during treatment. A mathematical model of these treatment dynamics confirmed that this wound-healing response could contribute to the amplification of the CSC population during chemotherapy [11]. At the same time, however, the model showed that this CSC enrichment was not maintained following cessation of therapy. Instead, CSC fractions were predicted to return to pre-treatment levels after treatment cessation. In other words, the positive feedback wound-healing response alone could not account for a pronounced reduction in the response to chemotherapy during the next cycle following an off-therapy phase. This situation changed when we added negative feedback from differentiated cells onto the division rate of stem cells in the mathematical model [11]. We found that in this case, the enriched stem cell fractions were predicted to be maintained in the long term during the off-treatment phase. The model with the negative feedback loop could thus reproduce the experimentally observed result that CSC enrichment during a round of chemotherapy could reduce the response to a subsequent chemotherapy cycle.

The notion that regulatory feedback loops originating from healthy tissue remain partially active in tumors is not well-established, nor is the notion that they determine the disease course or the response to therapy. The PGE_2_-based wound healing response in bladder cancer xeografts is the best experimental support for these ideas. Negative feedback loops have so far not been implicated in a similar way, but there is clear evidence that negative feedback regulation likely plays and important role in healthy tissue dynamics, e.g., in the olfactory epithelium, where GDF11 and Activin βB negatively regulate self-renewal rates in progenitor and stem cells [12,13]. Some of this negative feedback regulation could thus very well continue to be operational to a certain extent in tumors, and indeed the analysis of tumor growth data has found patterns that are difficult to account for in the absence of negative feedback regulation [14,15].

While our previous modeling showed that one particular negative feedback loop could lead to sustained CSC enrichment following chemotherapy of bladder cancer xenografts, many other regulatory feedback loops can potentially be present within the tumor. Some of these might also be able to contribute to sustained CSC enrichment and to a loss of therapy response, while others might not contribute to this phenomenon. A targeted experimental search for feedback factors that promote loss of treatment responses in bladder cancer requires us to know which types of feedback mechanisms intrinsic to the cell population can in principle promote sustained CSC enrichment. The aim of this paper is to identify such candidate feedback loops, based on the analysis of mathematical models, thus providing a theoretical basis for the experimental identification of relevant, specific feedback molecules.

We focus on two basic scenarios. First, we assume that partial breakage of feedback regulation results in temporary cell growth towards a new equilibrium, characterized by an overall larger number of cells. This could correspond to a single step in step-wise tumor progression. We investigate the conditions required for the stem cell fraction to be larger at the new compared to the old equilibrium. In particular we study how remaining feedback loops determine the stem cell fraction. Second, we consider unbounded tumor growth and investigate how different feedback mechanisms that remain in a growing tumor cell population can determine whether or not CSC enrichment occurs during growth. Much of this work is done using ordinary differential equations. In the context of unbounded tumor growth, the effect of spatial growth patterns on stem cell enrichment is explored using a stochastic agent-based model and an analytical approximation.

The present study provides a solid theoretical basis for implicating the presence of feedback regulatory loops as a determinant of responses to cancer therapy. This adds to the mathematical literature quantifying the role of feedback regulation for tissue and tumor dynamics [11,13,15-24], and builds upon the wider mathematical literature concerned with the dynamics of hierarchically structured cell populations, e.g. [25-31] and stem cell fractions [32]. Two major approaches can be mentioned as most relevant in the present context. One approach uses spatial, agent-based or hybrid models to investigate SC dynamics, including questions of SC enrichment. Reference [32] provides an excellent review of the relevant literature. In particular, a versatile model of CSC and non-SCs has been developed [33,34], where the cellular expansion and competition dynamics could be explored. It was found that an increased proliferation capacity of non-stem cells results in encapsulation of SCs and tumor dormancy, while an increase in migration may lead to ``liberation” and expansion of SCs. In [34] it was demonstrated that the CSC fraction of a tumor population can vary by multiple orders of magnitude as a function of the generational life span of the non-stem cancer cells. In [27], the time-dependent SC fraction was studied in the context of a similar model, and it was further found that spontaneous cell death yields a higher SC fraction; Reference [35] studied dynamics and localization of SCs by using a hybrid model.

The second approach is non-spatial, where the dynamics of SCs and their differentiation are represented by using ODEs with or without feedback. For example, in [24], a systematic analysis of feedback is performed by using delay differential equations. Reference [8] considers a two-compartment, multi-clone model with a specific type of negative feedback on SC self-renewal, and shows that clonal evolution selects for fast reproducing and highly self-renewing cells at primary diagnosis, while relapse following therapy-induced remission is associated with highly self-renewing but slowly proliferating cells. A multicompartment model of differentiation was introduced axiomatically in [25,26,36] and the steady states were studied, both for general functional forms and for a particular case of regulatory feedback functions. In [37], a similar model was used to compare cellular properties of leukemic stem cells to those of their benign counterparts, by deriving conditions for expansion of malignant cell clones. Reference [23] investigated the impact of feedback loops and their breakage on cancer progression. While questions of SC enrichment were not directly addressed in the above studies, reference [38] presented a hierarchical model of cell growth and differentiation, and showed that in exponential growth (and in the absence of any control loops) the fraction of SCs reaches a steady state level, which is a decreasing function of the number of compartments and an increasing function of SC proliferation rate.

While a wealth of results of SC fraction dynamics have been obtained in the spatial (agent based and hybrid) models, analytical understanding or generalizations are more difficult in the context of these models; also feedback factors were not usually considered explicitly. On the other hand, a high level of analytical understanding has been reached in ODE models, but SC enrichment dynamics has not been specifically studied, except in the simpler cases in the absence of feedback loops. In the present study, we develop an axiomatic model of SC dynamics where feedback loops are included explicitly, and the functional form of the controls (which remains unknown) is kept as general as possible. We investigate conditions that lead to SC enrichment under different types of growth dynamics, and define what types of regulatory feedback loops can and cannot contribute to CSC enrichment, providing guidance to further experimental inquiries into specific signaling mechanisms.

### The basic mathematical modeling approach

An ordinary differential equation model has been used to describe tissue hierarchy dynamics in a healthy tissue [13,16], and the models presented here build on these approaches. While cell lineages consist of stem cells, transit amplifying cells, and terminally differentiated cells, our models make a simplification and take into account only stem cells (which encompass all the proliferating cells) and differentiated cells. Denoting stem cells (SC) by x and differentiated cells (DC) by y, the model is given by:

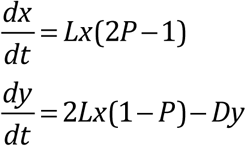

Stem cells divide with a rate *L*. With a probability *P*, the division results in two daughter stem cells (self-renewal), and with a probability (1-*P*), the division results in two daughter differentiated cells (differentiating division). Differentiated cells are assumed to die with a rate *D*. This model captures a probabilistic model of tissue control, which occurs on the population level if about half of the symmetric divisions result in two daughter stem cells and the other half result in two daughter differentiated cells. In addition to symmetric divisions, asymmetric divisions may play a role in tissue renewal. With asymmetric cell division, a stem cell gives rise to one stem cell and one differentiated cell, thus maintaining a constant population of stem cells. In the current model, even though such divisions do not appear explicitly, it is possible to show that they are included implicitly, see Section 1.2 of SI. It is proven that a model that considers both symmetric and asymmetric divisions is mathematically identical to the one studied here.

The basic model shown above contains no feedback loops. Hence, the rates *L* and *D* and probability *P* are constants that are independent of *x* or *y*. This system is only characterized by a neutrally stable family of nontrivial equilibria if *P*=0.5. If *P*>0.5, infinite growth is observed. If *P*<0.5, the cell population goes extinct.

It has been shown that introduction of negative feedback loops can result in more realistic behavior, where a stable equilibrium is attained for *P*>0.5 [13]. This was shown in the context of two specific feedback loops, and subsequently generalized to comprehensively list all possible (positive and negative) feedback loops compatible with stability [19,39,40]. Here, we also use a general model to assume different kinds of feedback on the rate of cell division, *L*, the rate of cell death, *D*, and the probability of self-renewal, *P*. We also add the possibility that stem cells die with a rate *δ* (which can also be subject to feedback). In the context of our model, feedback is equivalent to a dependence of rates and probabilities on the population sizes, *x* and/or *y*. Hence, the model is given by the following ODEs:

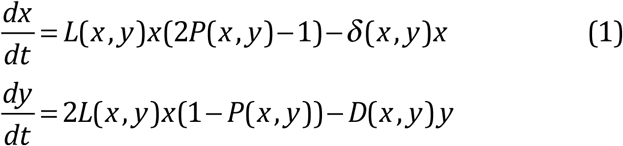

The division rate *L*, the death rates, *D* and *δ*, and the probability of self-renewal, *P*, are now functions of either the number of stem cells, *x*, or the number of differentiated cells, *y*, or both.

Evolution can result in the generation of mutant cell populations that are characterized by a higher self renewal probability, given by *P*_2_, which can be viewed as an early step towards carcinogenesis. Hence, we now have two stem and differentiated cell populations denoted by subscripts 1 and 2 for wild type and mutant types, respectively. The equations are thus given by

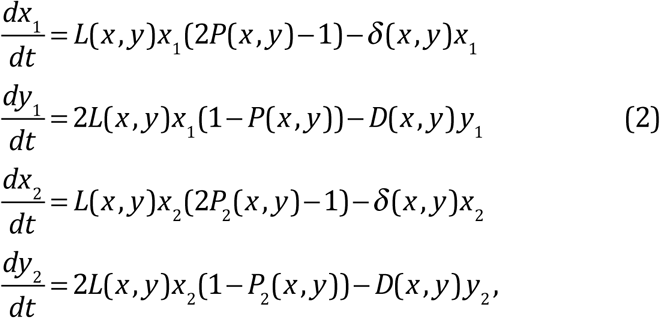

where *x=x*_*1*_*+x*_*2*_ and *y=y*_*1*_*+y*_*2*_. The two cell populations are in competition with each other, mediated by the feedback factors that are shared between the two populations. Table 1 summarizes all the variables used in this paper (both in this section and in the later sections). These equations are structurally similar to previously published models that were explored in different contexts [37].

**Table 1.**
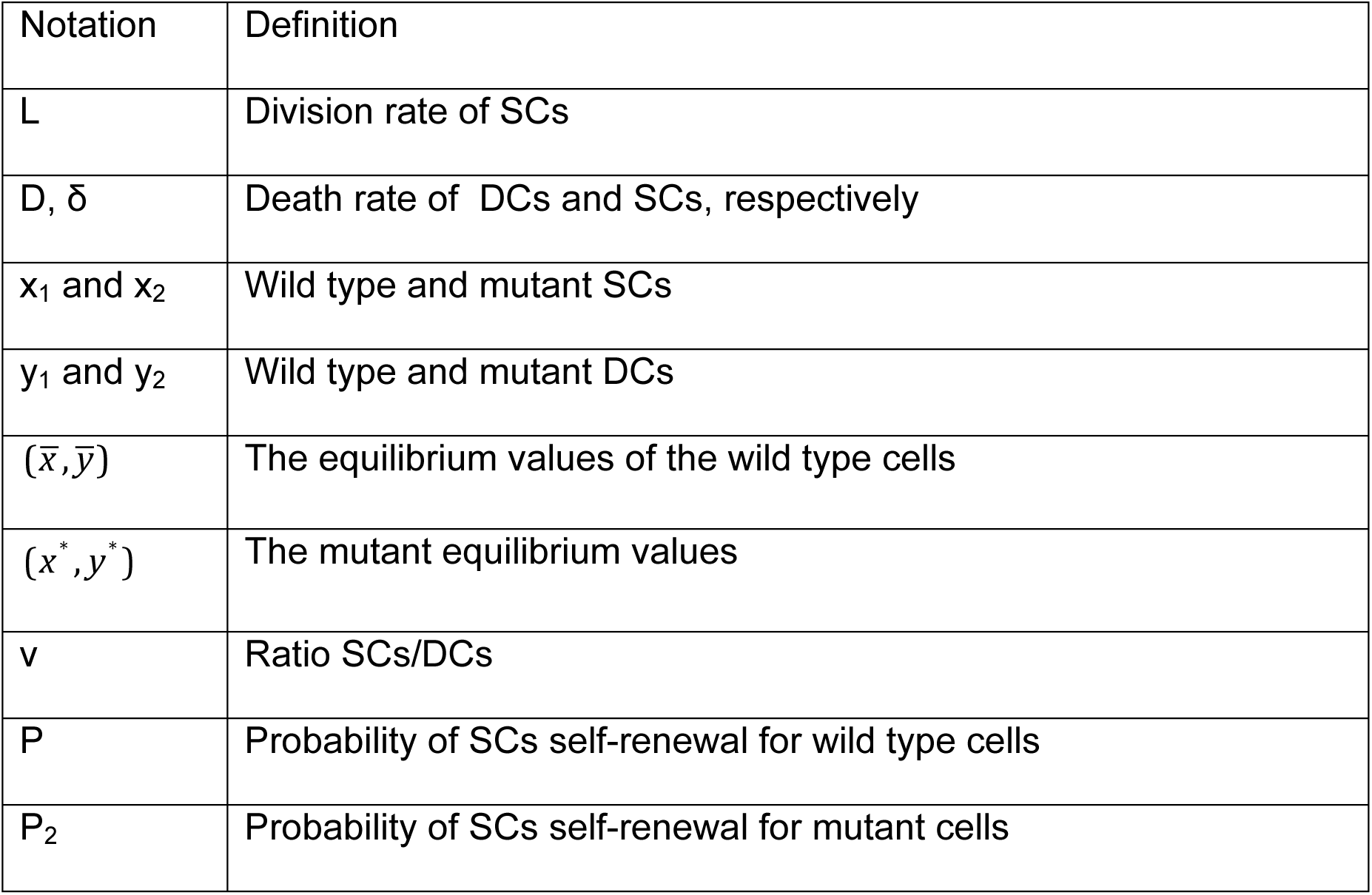
Mathematical symbols and the definitions of variables.

Note that we only consider mutants with a higher self-renewal probability, because these are the only mutant types that can grow from low numbers and invade in this type of model. Mutants in other parameters fail to grow and will die out, as described in a different context in reference [21]. In principle, it is possible that a mutation causes a simultaneous change in more than one parameter, which, however, does not lead to a change in conclusion and is not pursued further.

### Cell growth towards a new equilibrium

The first important scenario happens when the mutant cell population gains a selective advantage, outcompetes the original, healthy cell population, and grows towards a new and higher equilibrium level. This is achieved by assuming that *P*_2_>*P*, along with other conditions on the rate functions that are specified in Section 1 of SI We examine the conditions under which the stem cell fraction at the new equilibrium is increased compared to that at the original equilibrium. In terms of the model notation, we are interested in the quantity *v(x,y) = x / y*, i.e. the ratio of stem to differentiated cells. We investigate how different combinations of feedback loops that remain in the mutant cell population impact stem cell enrichment. In the following we assume that the division and death rates in the above models are monotonic functions of the number of stem and/or differentiated cells. We further assume that in the absence of mutants, the system is at equilibrium, characterized by the pair 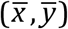, which satisfies:

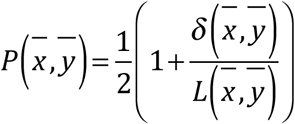

At this equilibrium, the fraction of stem to differentiated cells is given by

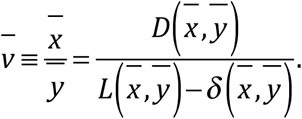

We assume that the mutant cell population can invade from low numbers and displace the original cell population. This occurs if *P*_2_(*x,y*) > *P*(*x,y*) (this is a sufficient condition). Assuming that a new equilibrium is reached, it is characterized

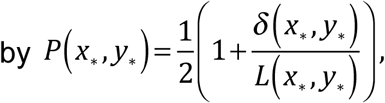

and the new fraction of stem cells is given by

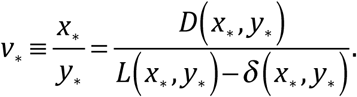

We examine under what conditions 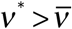, i.e. when the stem cell ratio at the newly obtained mutant equilibrium exceeds that at the original equilibrium.

We observe two qualitatively distinct outcomes, see Sections 2 and 3 of SI or the detailed analysis. (i) The fraction of stem cells increases compared to the previous equilibrium state. We call this stem cell enrichment. This occurs if the ratios *L/D* and/or *L/ δ* are decreasing functions of the cell population. This happens when larger cell population sizes result in negative feedback on cell expansion parameters and/or positive feedback on death. For example, if the per cell division rate decreases and/or the death rate increases with population size. (ii) The fraction of stem cells decreases compared to the previous equilibrium state. We call this stem cell depletion. This occurs if the ratios *L/D* or *L/ δ* are increasing functions of the cell population. In this scenario, larger populations promote cell expansion kinetics, e.g. by decreasing the death rate of cells or increasing their division rate. This would correspond to a positive feedback loop on cell division and/or a negative feedback loop on death rate. We note that mutant emergence generally changes the stem cell fraction, unless the division and death rates (*L, D*, and *δ*) are constant and hence not affected by feedback.

We will illustrate these points by using some specific examples. The first example is of SC enrichment. Consider a system where *δ*=0, the self renewal probabilities *P* and *P*_2_ are given by decreasing functions of y (see solid lines in Fig 1(a)), and the division rate *L* is also a function of y, solid line in Fig 1(b). The cell dynamics and control loops corresponding to this system are schematically shown in Fig 2(a). In this and other such diagrams, blue arrows correspond to cellular processes (characterized by kinetic rates, such as *L* and *D*), and cell fate decisions, which are probabilities (*P* or *P*_2_). The red negative arrows originating in the DC circle represent a negative dependence of both functions *L* and *P* on *y* (the number of DCs). In this first example, SC enrichment is predicted to occur. Fig 2(b) shows the cell dynamics, once a mutant is introduced at 100 time units. While the wild type population goes extinct, the mutants rise to a new equilibrium characterized by a significantly higher ratio *x/y* compared to the original equilibrium.

**Fig 1:**
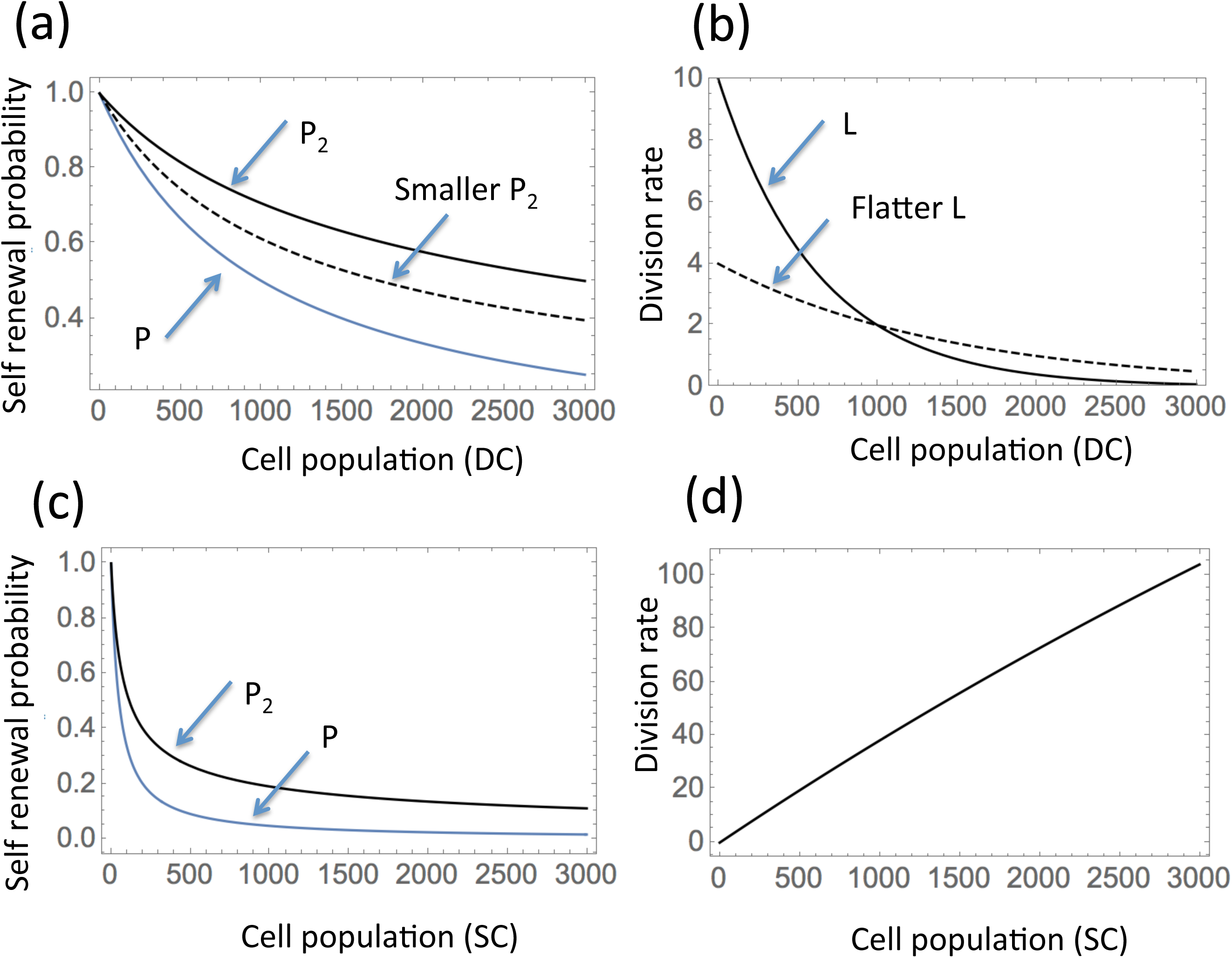
Examples of functional dependencies leading to SC enrichment (a,b) and depletion (c,d) at a new equilibrium. (a) Self renewal probability functions, *P*(*y*)=(1+0.001 *y*)^-1^ and *P* (*y*)=(1+0.001 *y*)^1/2^*P*(*y*) (the solid lines). An example of a smaller self renewal probability of mutants is given by *P* (*y*)=(1+0.005*y*) ^1/2^ *P*(*y*) (the dashed line). (b) An example of a negatively controlled division rate is given by *L*(*y*)=2×5^1-0.001y^ (solid line). A flatter division rate is given by *L*(*y*)=2^2-0.001y^ (dashed line). (c) Self renewal probability functions, *P*(*x*)=(1+0.02 *x*)^-1^ and *P* (*x*)=(1+0.015 *x*) ^1/2^ *P*(*x*). (d) An example of a positively controlled division rate is given by *L*(*x*)=401(1-e^-0.0001*x*^).

**Fig 2:**
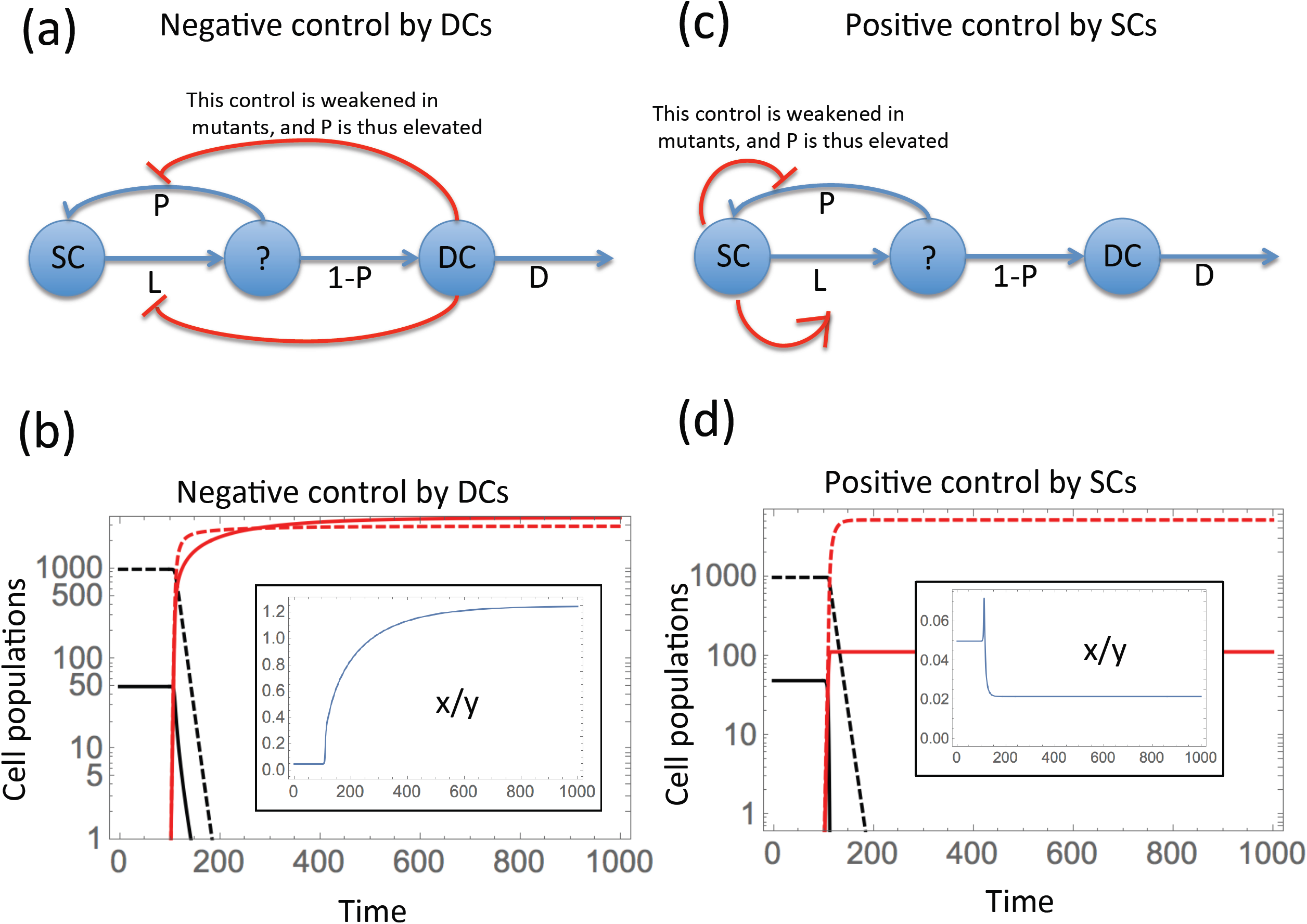
Dynamics of enrichment when a new equilibrium is reached by mutants. The left panels (a, b) illustrate SC enrichment, and the right panels (c,d) SC depletion. In the top panels, the dynamics and the control loops are presented schematically. SCs (the leftmost circle) divide at rate *L*, such that the decision of whether to self renew or to differentiate (denoted by a question mark) is governed by probability *P.* Control loops are depicted by red arrows (positive or negative) directed from the population mediating the control to the rate/probability that is being controlled. In the bottom panels, the time series are shown, where functions for wild type cells, *x*_*1*_ *(t)* and *y*_*1*_ *(t)*, are plotted in black, and functions for mutant cells, *x* _2_*(t)* and *y* _2_*(t)*, are plotted in red. SCs are depicted by solid and DCs by dashed lines. Initially, the system is at the original equilibrium, 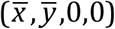. At t=100, mutant stem cells are introduced at a low level, resulting in the extinction of wild type cells, and convergence to a new equilibrium, (0,0,x*,y*). In the insets, the SC fraction, *x/y*, is depicted as a function of time. (a,b) Negative control by DCs, resulting in SC enrichment: the functions *L, P*, and *P*_*2*_ are given by the solid lines in Fig 1(a,b). (c,d) Positive control by SCs, resulting in SC depletion: the functions *L, P*, and *P*_*2*_ are specified in Fig 1(c,d).

The second example is SC depletion. It is given by a system with rate functions given by Fig 1(c,d), with δ=0 and the division rate positively controlled by the SC population. The controls are schematically shown in Fig 2(c). The resulting dynamics are presented in Fig 2(d), where the proportion of SCs at the new, mutant equilibrium is smaller than the original proportion. Note however that the dependence of *x/y* on time is non-monotonic and a temporary phase of SC enrichment is experienced before the ratio *x/y* lowers to its long-term level.

These two examples illustrated in Fig 2 show that the proportion of SCs can either increase (SC enrichment) or decrease (SC depletion), once mutants with altered (increased) self-renewal probability take over the cell population. We refer to Section 2 of SI for the detailed analysis of the scenario where all the rates and functions are controlled either by SCs or by DCs, but not both. Section 3 of SI extends this to the more general case where the rate functions depend both on *x* and *y*.

Next, we examine the factors that affect the magnitude of the change. In Fig 3 we take the basic model of Fig 2(a) and modify it in three different ways, each of which leads to a decrease in the amount of SC enrichment, compared to the situation in Fig 2(a). First, we consider the death rate of the stem cells, δ. The amount of enrichment is smaller for nonzero stem cell death rates, compared to the case of *δ*=0. In Fig 3(b) we can see that in the presence of SC death, the increase in the enrichment parameter, *x/y*, is more modest than that of Fig 2(b).

**Fig 3.**
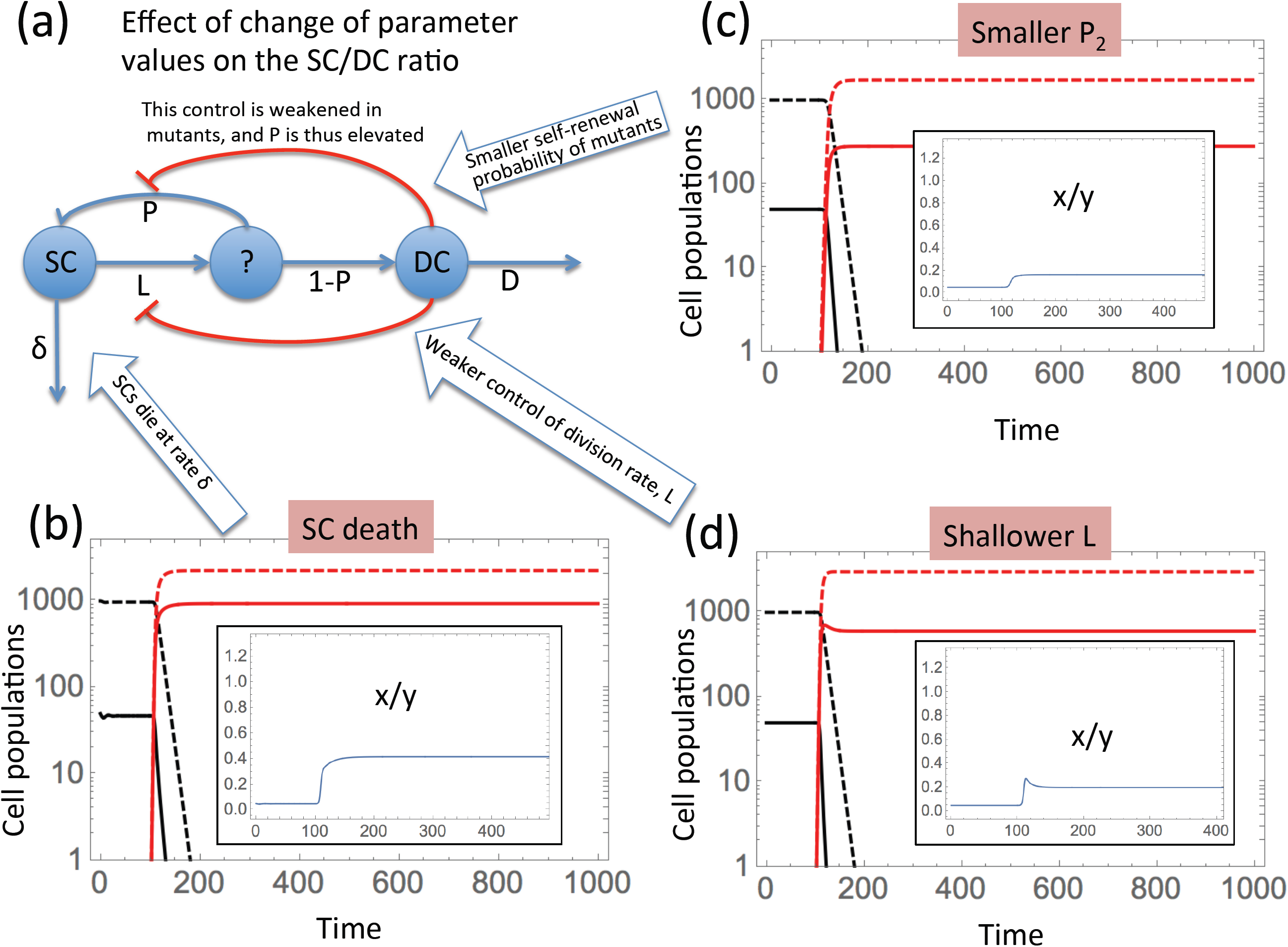
Parameter dependence of the SC enrichment magnitude. (a) The basic model, whose 3 aspects are modified. SCs (the leftmost circle) divide at rate *L*, such that the decision of whether to self renew or to differentiate (denoted by a question mark) is governed by probability *P*. Control loops are depicted by red arrows (positive or negative) directed from the population mediating the control to the rate/probability that is being controlled. (b) SC death: same parameters as in Fig 2(a), except *δ*=0.03. (c) Smaller *P*_*2*_: same parameters as in Fig 2(b) except *P*_*2*_ is given by the dashed line in Fig 1(a). (d) Shallower *L*: same parameters as in Fig 2(b), except the division rate *L* is given by the dashed line in Fig 1(b). We observe that the SC enrichment is (b-d) is less pronounced compared to Fig 2(b).

The second modification is a smaller self-renewal probability, *P*_2_, of the mutant cell population, Fig. 3(c). The self-renewal probability *P*_2_ of the mutant cells is that given by the dashed line in Fig 1(a). Again, this results in a more modest increase in the stem cell fraction compared to Fig 2(b).

The third modification is a less pronounced feedback on the division rate. The result is that the SC enrichment becomes smaller. In Fig 3(d) we use a flatter division rate function, *L*, than that in Fig 2(a) (compare the dashed line in Fig 1(b) with the solid line). This results in a smaller SC enrichment, as shown by a smaller increase of *x/y* in Fig 3(d) compared to Fig 2(b). A detailed analysis of all these scenarios is presented in Section 2 of SI.

### Stem cell enrichment in non-equilibrium situations

Next, we study the scenarios where the mutant population grows from low numbers and does not reach a new equilibrium, but instead, continues to grow indefinitely. This would correspond to tumor growth. The definition of enrichment in this context and the relevant methodology will be somewhat different in this case. We will consider the mutant population alone, and study the growth of *x*_2_ and *y*_2_, in order to find the dynamics of the quantity v=*x*_2_/*y*_2_. We will say that stem cell enrichment occurs if the quantity v increases during population growth, either infinitely, or temporarily. For simplicity we will assume that the cancer stem cell (CSC) population does not die (*δ*=0). We further assume that the probability of self-renewal of mutants, *P*_2_, is a monotonic function of the population size (stem cells or differentiated cells) that satisfies *P*_2_ > 1/2, and that in the limit of large populations, it approaches a limiting value, 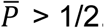 When we consider the mutant dynamics, we will drop the subscript 2, and simply study the equations:

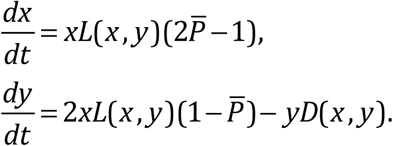

If we assume that both *D* and *L* depend on a single population, that is, *L=L(x), D=D(x)*, or *L=L(y),D=D(y)*, the following approximations can be derived (see Section 4 of SI). As the cell population expands, the DC population behaves as

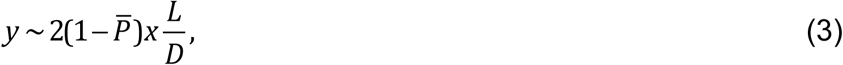

and ratio of stem to differentiated cells, v, in the limit of large times is given by:

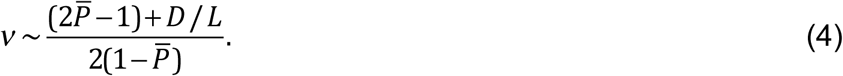

Equation (4) states that the long-term dynamics of the SC fraction, v, is defined by the behavior of the ratio *D/L*, the per cell death rate of the DCs and per cell division rate of SCs. If either or both of these quantities are controlled by the population size, the ratio will change as the tumor grows. If the behavior of the death rate and division rates are known, formula (4) predicts the dynamics of the SC fraction in tumor growth. Three scenarios are possible.

#### 1) Unlimited SC enrichment

One relevant scenario occurs if the ratio *D/L* grows indefinitely as the population size increases; this corresponds to either a negative feedback on *L* (such that *L* approaches zero as population size increases), and/or a positive feedback on *D*. If an increase in cell population sizes results in an unbounded increase in *D/L*, then the stem cell fraction, v, will continuously rise towards infinity. Therefore, as the tumor cell population grows, the fraction of SCs increases, and at very large tumor sizes, the tumor is practically only made up of CSCs. This scenario is illustrated in Figs 4(a,b) and 5(a,b), where negative feedback on *L* results in an increase in the ratio *D/L*, mediated by differentiated cells (panels (a)) and stem cells (panels (b)). Figs 4(a), 5(a) explore negative feedback on *L* by DCs. It depicts a system similar to that of Fig 2(a), except the self-renewal probability of the mutants is a constant, resulting in an unlimited growth of mutants. The fraction *x/y* in this case experiences unlimited growth, as shown by the inset in panel 5(a). In Figs 4(b) and 5(b), we replace control of *L* by DCs with control by SCs; again, an unlimited growth of *x/y* is observed, see the inset in panel 5(b).

**Fig 4.**
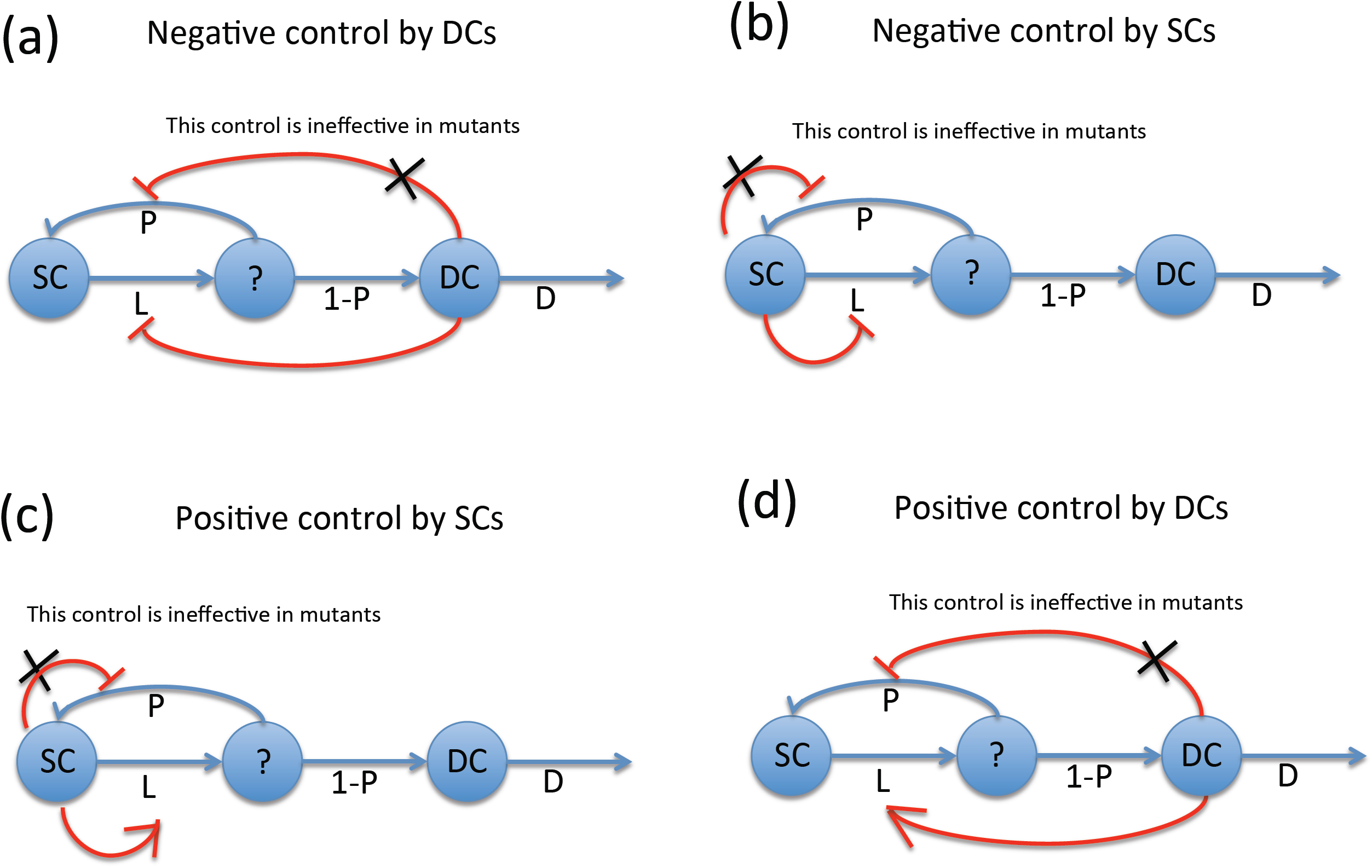
Four types of control used in Fig 5 to study unbounded growth. SCs (the leftmost circle) divide at rate *L*, such that the decision of whether to self renew or to differentiate (denoted by a question mark) is governed by probability *P*. Control loops are depicted by red arrows (positive or negative) directed from the population mediating the control to the rate/probability that is being controlled. (a) Negative control by DCs; (b) Negative control by SCs; (c) Positive control by DCs; Positive control by SCs; (d) Positive control by SCs. Unlike in Fig 2(a,c), the mutant self-renewal probability is assumed to be constant (and thus the control loop is completely severed in mutants), leading to unlimited growth. The 2 top panels correspond to Fig 5(a,b); the 2 bottom panels correspond to Fig 6(a,b).

**Fig 5.**
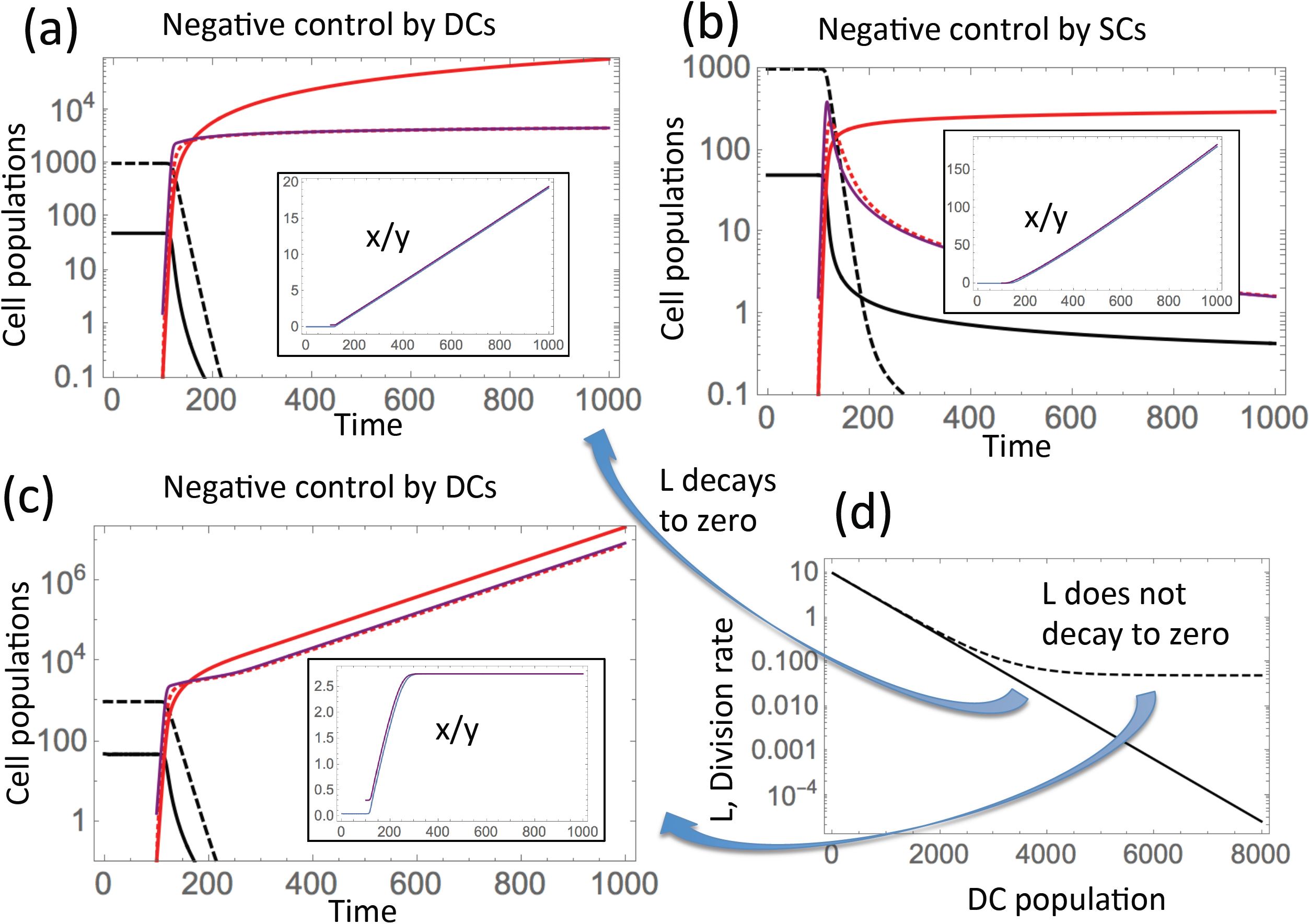
Dynamics of unbounded growth under negative control on SC divisions. Notations are as in Fig 2(b,d). The purple line in each panel plots the approximation for y (formula (3)), and in the insets the approximation for *x/y* (formula (4)). (a) Negative control by DCs: parameters are as in Fig 2(b) except 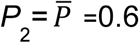. (b) Negative control by SCs: similar to (a), except the rate functions depend on SCs: *L(x)*=2*5^(1-*x*/50)-1^, *P(x)*=(1+0.02*x*)^-1^, and 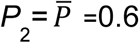. In both panes, unlimited SC enrichment is observed. (c) Saturated SC enrichment: same as (a), except the division rate decreases to a constant: *L(y)=*0.05+2*5^1-0.001*y*^. (d) The two division rates are plotted on a logarithmic scale as functions of DC populations. The solid line depicts the function used in panel (a) and the dashed line the function used in panel (c).

#### 2) Saturated SC enrichment

The second type of behavior is observed if an increase in the number of cells results in a saturated growth in *D* and/or a decay of *L* to a nonzero level, such that the ratio *D/L* is a growing and saturating function as the cell population increases. In this case, the ratio v will increase and reach a constant level, as the tumor size grows beyond a given size. In other words, once the tumor size has reached a critical threshold, the stem cell fraction is predicted to remain constant as the tumor grows further. This scenario is illustrated in figure 5(c,d), where populations x and y grow indefinitely, but the ratio x/y first increases and then reaches a constant level (see the inset). The only difference in simulations of panel 5(c) compared to those of panel 5(a) is the fact that the division rate, *L*, no longer decreases to zero, but reaches a small but nonzero constant level, as the cell population grows. Panel 5(d) depicts the two functions, *L(y)*, that are used in the simulation (the solid line for panel 5(a) and the saturating, dashed line for panel 5(c)).

#### 3) SC depletion

The third and final scenario that is possible corresponds to the case where function *D/L* decreases as the population size increases. This can happen if the division rate is positively controlled and/or when the death rate is negatively controlled by the cell populations. This type of dynamics is illustrated in figures Fig 4(c, d) and 6(a,b). In these simulations, positive feedback on *L* was mediated either by SCs (figures 4(c) and 6(a)) or by differentiated cells (figures 4(d) and 6(b)). In this case, a temporary reduction in the ratio v can be observed during tumor growth, until v converges to a constant. In other words, we observe a reduction in the CSC fraction over time, until it reaches a limiting value. Note that the SC fraction can never decrease to zero, the minimum fraction is given by 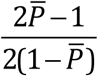.

**Fig 6.**
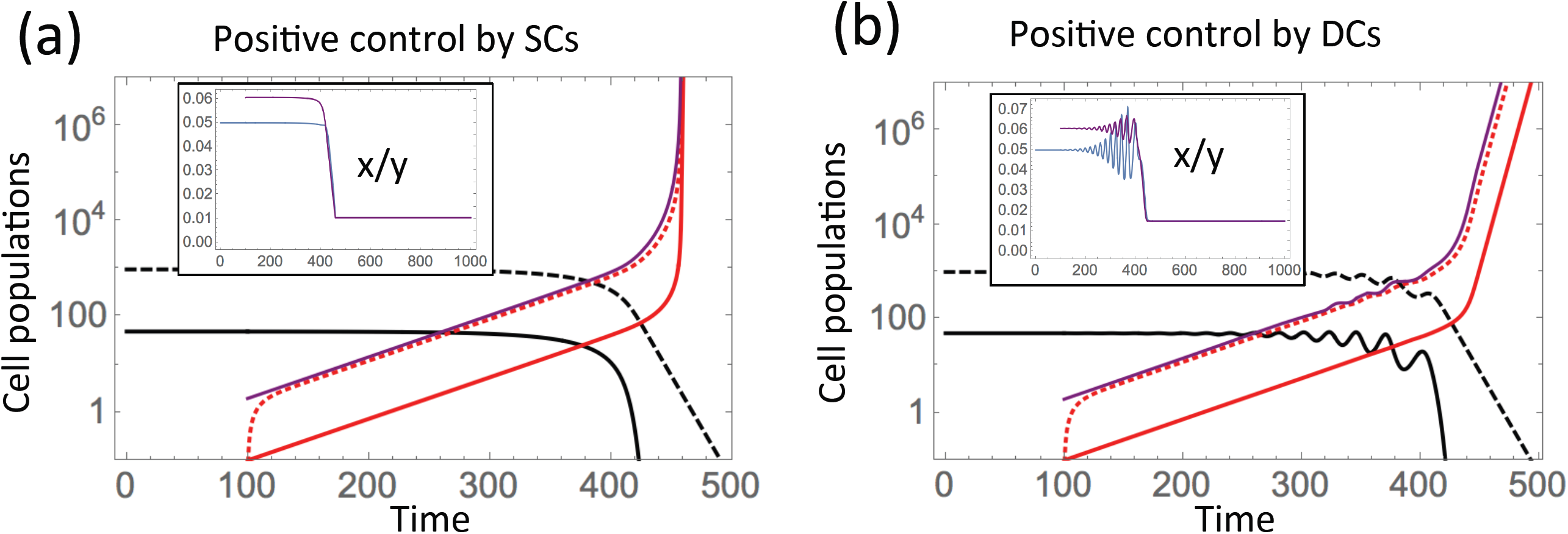
Dynamics of unbounded growth under positive control on SC divisions. Notations are as in Fig 2(b,d). The purple line in each panel plots the approximation for y (formula (3)), and in the insets the approximation for *x/y* (formula (4)). (a) Positive control by SCs: corresponds to parameters of Fig 2(d), except 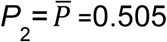. (b) Positive control by DCs: similar to (a), except the rate functions depend on DCs: *L*(y)=21(1-e^-0.0001*y*^), *P*(y)=(1+0.001*y*) ^-1^, 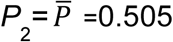.

To summarize, the following three types of SC fraction dynamics are predicted: (1) If *L* is subject to negative feedback and decays to zero and/or *D* is subject to positive feedback and increases without bound, then we have unlimited SC enrichment, such that the content of SCs grows to 100% in the log run. (2) If *L* is subject to negative feedback but never decreases to zero and/or *D* is subject to positive feedback and increases within bounds, then we have limited SC enrichment, where the SC fraction increases and then remains constant. (3) If *L* is subject to positive feedback and/or *D* is subject to negative feedback, then we have SC depletion, and the SC fraction decreases to a nonzero level.

Note that if *L* and *D* are subject to the same type of feedback, then the resulting behavior would be determined by the overall behavior of *L/D*. For example, if *L* and *D* are both subject to positive feedback and increase within bounds, but the feedback on *L* is greater than the feedback on *D* such that *L/D* is increasing with the population size, then this would correspond to type (3) and result in depletion. However, if the feedback on *D* is (effectively) greater such that *L/D* is decreasing and bounded above 0, then this would correspond to type (2) and result in limited SC enrichment, where SC fraction grows and saturates below 100%.

### SC enrichment in spatially structured populations

According to the above results, stem cell enrichment during growth requires certain feedback mechanism to be present in the tumor cell population, such that the ratio *L/D* is reduced as the population size is increased. Typically such feedback can occur through signaling factors that are secreted from stem or differentiated cells. Another way to achieve a similar result can be spatially restricted reproduction of cells. In such scenarios, cells experience range expansion in two dimensions, or grow as expanding sphere-like structures in 3D. Inside the expanding population, divisions must be balanced with deaths because free space is limited. In a way this works similarly to control loops affecting division and/or death rates, which were discussed earlier in the paper; in the case of spatially restricted growth, control is essentially competition for space, which leads to slower divisions/higher death as the density increases.

To explore this, we first considered a two-dimensional stochastic agent-based model that describes spatially restricted cell growth. The model assumes a 2-dimensional grid consisting of *n*x*n* spots. A spot can either be empty, contain a stem cell, or contain a differentiated cell. At each time step, the grid is sampled *N* times, where *N* is the number of cells currently present in the grid. If the sampled spot contains a stem cell, it divides with a probability *L*_0_, and dies with a probability *δ*_0_. If the division event is chosen, one of the eight nearest neighboring spots is randomly picked as a target for one of the daughter cells. If the chosen spot is already filled, the division event is aborted, otherwise it proceeds. If division proceeds, both daughter cells will be stem cells with a probability *P*_0_ (self-renewal). With probability 1-*P*_0_, both daughter cells will be differentiated cells. If the sampled spot contains a differentiated cell, death occurs with a probability *D*_0_. No explicit feedback processes were included in the model. The simulation was started with 9 stem cells (and no differentiated cells). Assuming that stem cells do not die, the resulting average growth curve is shown in Figure 7(a). While initially, the differentiated cells grow to be more abundant than the stem cells, the stem cell population enriches over time and eventually becomes dominant as the cell population grows. Figure 7(b) shows the same kind of simulation, but assuming that stem cells die with a rate that is smaller than the death rate of differentiated cells. Consistent with the results obtained for explicit feedback mechanisms, we find that the degree of stem cell enrichment is reduced in the presence of stem cell death (higher rates of stem cell death lead to less enrichment).

**Figure 7.**
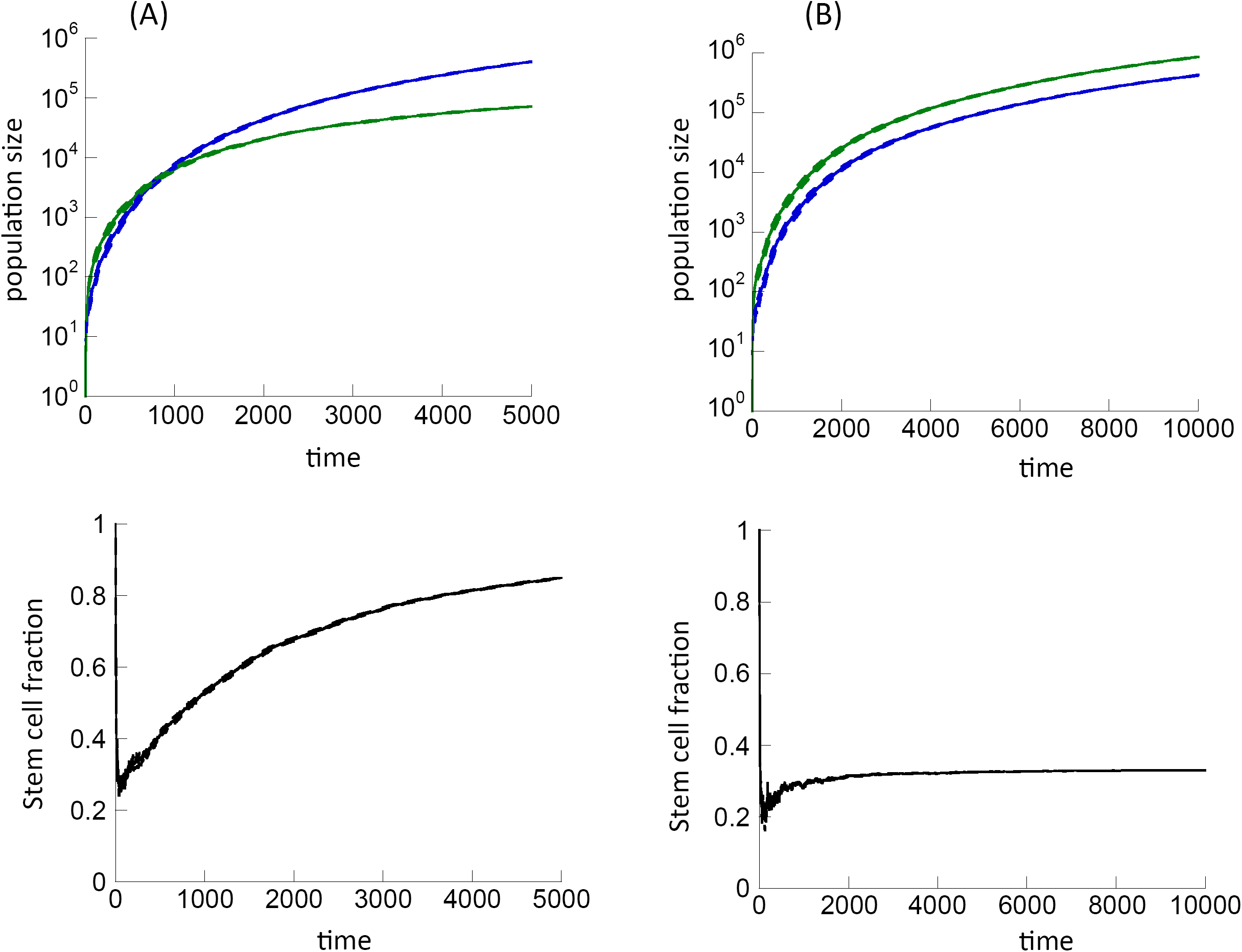
Effect of spatially restricted cell division on CSC enrichment, according to the 2-dimensional implementation of the agent-based model. (A) Dynamics in the absence of stem cell death, *δ*_0_=0. In the top panel, stem cells are shown in blue, differentiated cells in green. The bottom panel shows the stem cell fraction over time. (B) Same, but for a non-zero stem cell death rate, i.e. *δ*_0_=0.005. Other parameters were chosen as follows: *L*_0_=0.95, *P*_0_=0.6, *D*_0_=0.01, *n*x*n*=1500^2^. 8 simulations were run, and the mean is shown as a solid line while the mean ± standard error are shown by dashed lines.

A simple mean-field model that takes account of the space limitations inside an expanding population can explain these results (see Section 5 of SI). Indeed, the system in the interim of the expanding globe reaches a dynamic equilibrium state where the density of SCs and DCs is dictated by the balance of division and death rates. Figure 8 shows theoretical predictions for the densities of SCs and the DCs in the colony’s interim; the solid lines depict theoretical predictions and points correspond to numerical simulations of the agent based model. The figure also provides typical images for two parameter combinations; blue dots represent SCs and yellow dots DCs. As expected, under higher DC death the proportion of DCs decreases.

**Figure 8.**
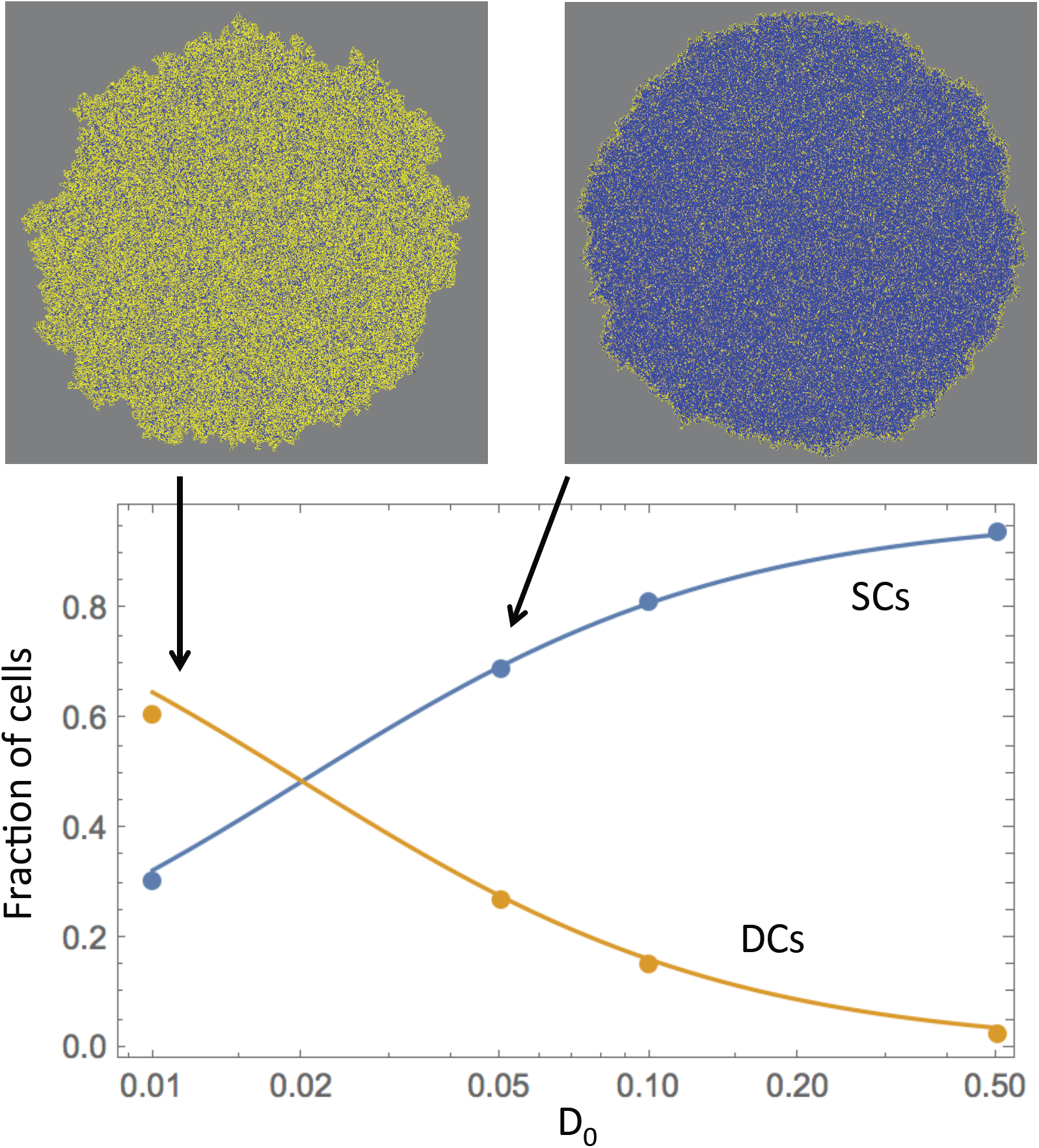
Agent based model simulations and theory. The fractions of SCs and DCs are shown for different values of parameter *D*_0_. Theoretical predictions from mean field modeling are given by solid lines and agent based model simulation results by points. Typical simulation results are presented for two of the parameter combinations. Gray is empty spots, blue SCs and yellow DCs. The rest of the parameters are: *L*_0_=0.95, *P*_0_=0.6, *δ*_0_=0.005.

Using the mean-field model, we have also calculated the degree of SC enrichment. The higher the SC death rate, the lower the overall density and the higher the fraction of DCs in the population. As time increases, the fraction of SCs reaches the value given by

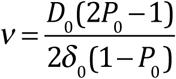

In particular, in the absence of SC death (*δ*_0_=0), the equilibrium value of v is infinity, which means that SCs exclude DCs everywhere in the core of the expanding range, such that DCs only concentrate in the exterior part of the colony. This corresponds to unlimited SC enrichment.

Agent-based simulations were repeated assuming a 3-dimensional space, where offspring cells could be placed into one of the 27 nearest neighboring spots (Section 5 of SI). Because there are more neighbors in three dimensions, the system is more mixed, and the extent of stem cell enrichment is generally lower. (As mentioned in the beginning of this paper, no stem cell enrichment occurs in a perfectly mixed system in the absence of explicit feedback mechanisms).

## Discussion and Conclusions

The experimental observation that a wound-healing type mechanism in bladder cancer can modulate the responsiveness of the tumor to chemotherapy [10] is to our knowledge the first clear case where a specific feedback loop has been implicated in determining therapy outcome. At the same time, however, this interesting observation has brought up a number of questions that can best be answered by using mathematical models to interpret the data. Mathematical modeling [11] suggested that the wound-healing response alone cannot result in CSC enrichment that is sustained beyond the treatment phase, and hence cannot explain the loss of therapy response during subsequent treatment cycles. The mathematical model predicted, however, that inclusion of a particular negative feedback loop from differentiated cells onto the rate of CSC division can result in sustained CSC enrichment and in a loss of therapy response in successive treatment cycles. Many different negative and positive feedback loops, however, might be operational in bladder cancer, and so far we did not know which types of feedback mechanisms can in principle contribute sustained CSC enrichment (and a loss of treatment response), and which types cannot contribute to this phenomenon. This limits the experimental search for particular feedback factors that drive the response of bladder cancer to chemotherapy. The analysis presented here uses mathematical approaches to clearly define which types of feedback loops can and cannot contribute to sustained CSC enrichment, and this forms an important guide for the search of specific feedback factors in bladder cancer, and also potentially in other tumors in which CSC-based resistance turns out to be important.

We found that whether or not stem cell enrichment is observed in the system depends on how the ratio *D/L* (deaths to divisions) changes with the population size. If it rises as the population size increases, which corresponds to either a negative feedback on *L* or a positive feedback on *D*, exerted either by stem or by differentiated cells, then SC enrichment is observed. On the other hand, if the ratio *D/L* decreases as the population grows, then no stem cell enrichment is predicted, and in fact SC depletion can be observed. A simple analytical approximation is derived for the long-term behavior of the expected fraction of SCs in the population, which is novel and useful for data interpretation. Further, according to the model, three separate factors can decrease and even altogether prevent SC enrichment. One is the presence of SC death; the second is a very weak feedback (or no feedback) on SC divisions remaining in the mutant populations, and the third is a relatively low self-renewal probability of mutant cells.

In contrast to previous approaches, we employed axiomatic modeling techniques, where model parameters such as the division rate, death rate, and self-renewal probability of cells were general functions of the number of stem and differentiated cells, which can potentially secrete feedback factors. We have thus considered a class of mathematical models that incorporate a large variety of possible feedback loops that regulate cell fate decisions in lineages. The models have the flexibility as to which cell populations exert the feedback signals, and whether feedback loops are positive or negative. The (unknown) exact details of the feedback functions remain unspecified. This is an important innovation when mathematically modeling complex stem cell dynamics with feedback regulation. When considering a specific model, as has been done in much of the literature so far, results can depend on the particular mathematical formulation of feedback interactions. These particular formulations are typically arbitrary, and the same biological processes can often be described by somewhat different mathematical terms, which can impact model predictions and thus lead to robustness issues. The axiomatic modeling approach performed here does not suffer from this problem.

The spatial models considered here have shown that in the presence of spatially restricted cell division, CSC enrichment can occur even in the absence of explicit feedback loops mediated by signaling molecules. It is the nature of spatially restricted cell spread that cells in the inside of an expanding population compete for space, such that their reproduction rate declines with population size and/or their death rate increases. This has the same effect on the dynamics as the presence of explicit feedback loops. Whether this represents a physiologically important mechanism that drives CSC enrichment remains to be explored further. The presence of cell migration can destroy the reported effect (because cell migration essentially leads to a higher degree of cell mixing). Since cell migration occurs even in spatially structured tumors, it is likely that explicit feedback regulatory loops have to be invoked to explain the loss of treatment response in bladder cancer.

All modeling approaches contain simplifying assumptions, and the models presented here are no exception. To gain analytical insights, we reduced the complexity of the lineage differentiation pathway to include only stem cells and differentiated cells, ignoring intermediate transit amplifying cell populations with limited self-renewal capacity. Our previous work included models that explicitly took into account transit amplifying cells [11], and the relationship between the presence of negative feedback on stem cell division and the occurrence of stem cell enrichment remained qualitatively the same [41]. Other modeling approaches have treated the cell differentiation pathway as a continuous process using partial differential equations, rather than considering discrete cell sub-populations [42]. Future work is required to determine to what extent the dynamics explored here remain robust in those types of models.

It is further important to note that the models discussed here are based on a set of core assumptions that we consider critical when addressing the particular questions that were the focus of this investigation. Further complexities can be included in future work when studying more advanced questions. An important example is additional tumor heterogeneity that is certainly present in bladder caners as well as in most other tumors. Variation is likely to occur in division rates, death rates, and the ability to secrete or respond to feedback signals; it will be important to extend the models with this in mind. Furthermore, while we concentrated on feedback signals that originate from tumor cells themselves, it is likely that regulatory signals originating from the microenvironment play a role in shaping cell renewal and differentiation dynamics. Thus, there is indication that in bladder cancer, interactions between the tumor cells and the extracellular matrix are important for pathogenesis and treatment [43]. These interactions are worth exploring mathematically in their own right, and would go beyond the scope of the current paper.

## Acknowledgements

We would like to thank David Axelrod for valuable suggestions about the manuscript. We acknowledge partial funding from the National Institutes of Health (NIH) through grant 1U54CA217378-01A1 for a National Center in Cancer Systems Biology at the University of California, Irvine, and NIH grant P30CA062203 for the Chao Comprehensive Cancer Center at the University of California, Irvine. In addition, NK and DW acknowledge NIH grant 1U01CA187956, and JL acknowledges partial support from the National Science Foundation, Division of Mathematical Sciences under grant NSF-DMS-1714973.

## References

1. Dean M, Fojo T, Bates S (2005) Tumour stem cells and drug resistance. Nat Rev Cancer 5: 275–284.

2. Brooks MD, Burness ML, Wicha MS (2015) Therapeutic Implications of Cellular Heterogeneity and Plasticity in Breast Cancer. Cell Stem Cell 17: 260–271.

3. Batlle E, Clevers H (2017) Cancer stem cells revisited. Nat Med 23: 1124–1134.

4. Shibue T, Weinberg RA (2017) EMT, CSCs, and drug resistance: the mechanistic link and clinical implications. Nat Rev Clin Oncol 14: 611–629.

5. Bao S, Wu Q, McLendon RE, Hao Y, Shi Q, et al. (2006) Glioma stem cells promote radioresistance by preferential activation of the DNA damage response. Nature 444: 756–760.

6. Diehn M, Cho RW, Lobo NA, Kalisky T, Dorie MJ, et al. (2009) Association of reactive oxygen species levels and radioresistance in cancer stem cells. Nature 458: 780–783.

7. Brock A, Chang H, Huang S (2009) Non-genetic heterogeneity--a mutation-independent driving force for the somatic evolution of tumours. Nat Rev Genet 10: 336–342.

8. Stiehl T, Baran N, Ho AD, Marciniak-Czochra A (2014) Clonal selection and therapy resistance in acute leukaemias: mathematical modelling explains different proliferation patterns at diagnosis and relapse. J R Soc Interface 11: 20140079.

9. Hamis S, Nithiarasu P, Powathil GG (2018) What does not kill a tumour may make it stronger: In silico insights into chemotherapeutic drug resistance. J Theor Biol 454: 253–267.

10. Kurtova AV, Xiao J, Mo Q, Pazhanisamy S, Krasnow R, et al. (2015) Blocking PGE2-induced tumour repopulation abrogates bladder cancer chemoresistance. Nature 517: 209–213.

11. Rodriguez-Brenes IA, Kurtova AV, Lin C, Lee YC, Xiao J, et al. (2017) Cellular Hierarchy as a Determinant of Tumor Sensitivity to Chemotherapy. Cancer Res 77: 2231–2241.

12. Gokoffski KK, Wu HH, Beites CL, Kim J, Kim EJ, et al. (2011) Activin and GDF11 collaborate in feedback control of neuroepithelial stem cell proliferation and fate. Development 138: 4131–4142.

13. Lander AD, Gokoffski KK, Wan FY, Nie Q, Calof AL (2009) Cell lineages and the logic of proliferative control. PLoS Biol 7: e15.

14. Rodriguez-Brenes IA, Komarova NL, Wodarz D (2013) Tumor growth dynamics: insights into evolutionary processes. Trends Ecol Evol 28: 597–604.

15. Youssefpour H, Li X, Lander AD, Lowengrub JS (2012) Multispecies model of cell lineages and feedback control in solid tumors. J Theor Biol 304: 39–59.

16. Kunche S, Yan H, Calof AL, Lowengrub JS, Lander AD (2016) Feedback, Lineages and Self-Organizing Morphogenesis. PLoS Comput Biol 12: e1004814.

17. Komarova NL, van den Driessche P (2018) Stability of Control Networks in Autonomous Homeostatic Regulation of Stem Cell Lineages. Bull Math Biol 80: 1345–1365.

18. Konstorum A, Hillen T, Lowengrub J (2016) Feedback Regulation in a Cancer Stem Cell Model can Cause an Allee Effect. Bull Math Biol 78: 754–785.

19. Komarova NL (2013) Principles of regulation of self-renewing cell lineages. PLoS One 8: e72847.

20. Yang J, Plikus MV, Komarova NL (2015) The Role of Symmetric Stem Cell Divisions in Tissue Homeostasis. PLoS Comput Biol 11: e1004629.

21. Rodriguez-Brenes IA, Komarova NL, Wodarz D (2011) Evolutionary dynamics of feedback escape and the development of stem-cell-driven cancers. Proc Natl Acad Sci U S A 108: 18983–18988.

22. Rodriguez-Brenes IA, Wodarz D, Komarova NL (2013) Stem cell control, oscillations, and tissue regeneration in spatial and non-spatial models. Front Oncol 3: 82.

23. Stiehl T, Ho AD, Marciniak-Czochra A (2018) Mathematical modeling of the impact of cytokine response of acute myeloid leukemia cells on patient prognosis. Sci Rep 8: 2809.

24. Arino O, Kimmel M (1986) STABILITY ANALYSIS OF MODELS OF CELL PRODUCTION SYSTEMS Mathematical Modeling 7: 1269–1300.

25. Marciniak-Czochra A, Stiehl T, Ho AD, Jager W, Wagner W (2009) Modeling of asymmetric cell division in hematopoietic stem cells--regulation of self-renewal is essential for efficient repopulation. Stem Cells Dev 18: 377–385.

26. Stiehl T, Marciniak-Czochra A (2011) Characterization of stem cells using mathematical models of multistage cell lineages. Mathematical and Computer Modelling https://doi.org/10.1016/j.mcm.2010.03.057.

27. Enderling H, Hlatky L, Hahnfeldt P (2013) Cancer Stem Cells: A Minor Cancer Subpopulation that Redefines Global Cancer Features. Front Oncol 3: 76.

28. Werner B, Dingli D, Lenaerts T, Pacheco JM, Traulsen A (2011) Dynamics of mutant cells in hierarchical organized tissues. PLoS Comput Biol 7: e1002290.

29. Michor F (2008) Mathematical models of cancer stem cells. J Clin Oncol 26: 2854–2861.

30. Glauche I, Cross M, Loeffler M, Roeder I (2007) Lineage specification of hematopoietic stem cells: mathematical modeling and biological implications. Stem Cells 25: 1791–1799.

31. Roeder I, Loeffler M (2002) A novel dynamic model of hematopoietic stem cell organization based on the concept of within-tissue plasticity. Exp Hematol 30: 853–861.

32. Enderling H (2014) Cancer stem cells: small subpopulation or evolving fraction? Integrative Biology 7: 14–23.

33. Enderling H, Anderson AR, Chaplain MA, Beheshti A, Hlatky L, et al. (2009) Paradoxical dependencies of tumor dormancy and progression on basic cell kinetics. Cancer Res 69: 8814–8821.

34. Enderling H, Hlatky L, Hahnfeldt P (2009) Migration rules: tumours are conglomerates of self-metastases. Br J Cancer 100: 1917–1925.

35. Sottoriva A, Verhoeff JJ, Borovski T, McWeeney SK, Naumov L, et al. (2010) Cancer stem cell tumor model reveals invasive morphology and increased phenotypical heterogeneity. Cancer Res 70: 46–56.

36. Nakata Y, Getto P, Marciniak-Czochra A, Alacron T (2012) Stability analysis of multi-compartment models for cell production systems. Journal of Biological Dynamics 6(Suppl. 1): 2–18.

37. Stiehl T, Marciniak-Czochra A (2012) Mathematical Modeling of Leukemogenesis and Cancer Stem Cell Dynamics. Mathematical Modelling of Natural Phenomena 7: 166–202.

38. Weekes SL, Barker B, Bober S, Cisneros K, Cline J, et al. (2014) A multicompartment mathematical model of cancer stem cell-driven tumor growth dynamics. Bull Math Biol 76: 1762–1782.

39. Komarova NL, van den Driessche P (2017) Stability of Control Networks in Autonomous Homeostatic Regulation of Stem Cell Lineages. Bull Math Biol.

40. Yang J, Axelrod DE, Komarova NL (2017) Determining the control networks regulating stem cell lineages in colonic crypts. J Theor Biol 429: 190–203.

41. Rodriguez-Brenes IA, Wodarz D, Komarova NL (2015) Characterizing inhibited tumor growth in stem-cell-driven non-spatial cancers. Math Biosci 270: 135–141.

42. Doumic C, Marciniak-Czochra A, Perthame B, Zubelli JP A structured population model of cell differentiation. SIAM Journal of Applied Mathematics 71: 1918–1940.

43. Brooks M, Mo Q, Krasnow R, Ho PL, Lee YC, et al. (2016) Positive association of collagen type I with non-muscle invasive bladder cancer progression. Oncotarget 7: 82609–82619.

